# Noise-guided tuning of synthetic protein waves in living cells

**DOI:** 10.1101/2025.03.21.644572

**Authors:** Dennis T. Bolshakov, Elliott W. Z. Weix, Thomas M. Galateo, Rohith Rajasekaran, Scott M. Coyle

## Abstract

Biological systems use protein circuits to organize cellular activities in space and time, but engineering synthetic dynamics is challenging due to stochastic effects of genetic and biochemical variation on circuit behavior. Genetically encoded oscillators (GEOs) built from bacterial MinDE-family ATPase and Activator modules generate fast orthogonal protein waves in eukaryotic cells, providing an experimental model system for genetic and biochemical coordination of synthetic protein dynamics. Here, we use budding yeast to experimentally define and model phase portraits that reveal how the breadth of frequencies and amplitudes available to a GEO are genetically controlled by ATPase and Activator expression levels and noise. GEO amplitude is encoded by ATPase absolute abundance, making it sensitive to extrinsic noise on a population level. In contrast, GEO frequency is remarkably stable because it is controlled by the Activator:ATPase ratio and thus affected primarily by intrinsic noise. These features facilitate noise-guided design of different expression strategies that act as filters on GEO waveform, enabling us to construct clonal populations that oscillate at different frequencies as well as independently tune frequency and amplitude variation within a single population. By characterizing 169 biochemically distinct GEOs, we provide a rich assortment of phase portraits as starting points for application of our waveform engineering approach. Our findings suggest noise-guided design may be a valuable strategy for achieving precision control over dynamic protein circuits.

## Introduction

Cells use networks of interacting genes and proteins to build molecular circuits that organize and regulate their dynamic behavior.^1–3^ While genetic circuits can control expression over long timescales, protein circuits are essential to rapidly sense and manipulate biological activities with spatiotemporal precision.^4–6^ Protein circuits implement many key dynamic cellular tasks like motility,^7,8^ information processing,^9^ phagocytosis,^10^ polarized secretion,^11^ and force generation.^12^ As such, there is increasing interest in designing programmable protein circuits with fast predictable dynamics.^13–15^

A protein circuit’s dynamic behavior is shaped at two levels. At one level, the activities of the protein components and their regulatory relationships establish a network structure that defines how their spatiotemporal dynamics unfold. For example, a coherent-feed forward network structure is well-suited to implement delays, while a mutual inhibition architecture facilitates polarization.^16,17^ At a second level, protein circuits are affected by the concentrations of their components, which are controlled upstream by gene expression. Differences in expression levels can cause the same network architecture to produce different outcomes.^18^ As a result, engineering dynamic protein circuit behaviors can be challenging, as both the genetic and biochemical contributions to circuit performance must be carefully coordinated to realize a target behavior.

One source of expression variation in protein circuits arises from stochastic expression noise, which introduces uncertainty in the concentrations of circuit components of genetically identical cells.^19^ This noise can be partitioned into an extrinsic component, which reflects expression differences between individual cells in the population; and an intrinsic component, which reflects differences in expression from equivalent promoters within the same cell.^19^ These sources of noise create a distribution of component levels within the population, introducing cell-to-cell variation in circuit dynamics. Noise effects can be undesirable if a specific dynamic output is critical for function. Alternatively, structured variation can also be a feature that allows probabilistic behaviors to be encoded through a diversity of functional but quantitatively distinct outputs.^20^ The effects and consequences of noise have been explored for many natural endogenous signaling networks.^21,22^ Noise-guided strategies for synthetic protein circuit design could improve our ability to genetically encode specific dynamic outputs at a population level.

Recently, genetically encoded oscillators (GEOs) built from MinDE-family ATPase and Activator modules have been developed to generate fast orthogonal protein waves for spatiotemporal protein circuit design in eukaryotic cells.^23^ These waves propagate through a reaction-diffusion mechanism: ATP-dependent assembly of the ATPase on membranes stimulates recruitment of its associated Activator, stimulating ATP hydrolysis and membrane release.^24–30^ The resulting wave produces a robust oscillating signal in the cell whose frequency and amplitude can be quantified using Fast Fourier Transform (FFT).^23,31^ GEO circuits have demonstrated a range of biotechnology applications including frequency-domain single-cell barcoding, AM/FM broadcasting of single-cell state dynamics, and sub-cellular patterning of biological activities.^23,32^ The simplicity, orthogonality, and dynamic phenotypes of GEOs, combined with applications that rely on quantitative control of oscillations, makes them an ideal experimental testbed and model system for exploring genetic and biochemical coordination of synthetic protein dynamics.

Here, we implement GEOs in a budding yeast system to comprehensively characterize the genetic encodability of oscillations and engineer noise-guided circuit expression strategies that enable precise and tunable control over the resulting dynamics. We experimentally define and model phase portraits that reveal how the breadth of frequencies and amplitudes available to a GEO are genetically controlled by ATPase and Activator expression levels and noise. The GEO phase portrait reveals separable effects of extrinsic and intrinsic expression noise on GEO amplitude and frequency respectively. We exploit these differences to engineer different GEO expression strategies that act as noise filters on the phase portrait, allowing us to construct clonal populations that robustly oscillate at different frequencies or to independent tune frequency and amplitude spread within a single population. To expand GEO programmability, we characterized 169 biochemically distinct GEOs, providing a rich assortment of phase portraits spanning different frequency and amplitudes for waveform design. Our strategy of coordinating expression noise with circuit behavior may provide a more general strategy for improving precision control over protein dynamics in cells.

## Results

### Encodability of GEO protein oscillations is determined by phase portrait structure

Genetically encoded oscillators (GEOs) derived from bacterial MinDE-family ATPase/activator pairs robustly generate synthetic orthogonal protein waves *in vitro* and in eukaryotes.^23,27,28,32^ To systematically define the precision with which GEO oscillations can be genetically encoded and the impact of expression noise on oscillation behavior, we deployed GEOs in the budding yeast *Saccharomyces cerevisiae* (Fig. 1A). Budding yeast provide an ideal chassis for this inquiry, as a rich collection of expression vectors, promoter libraries, and single-copy integration strategies have been developed which allow for comprehensive manipulation of gene expression.^33^

**Figure 1:**
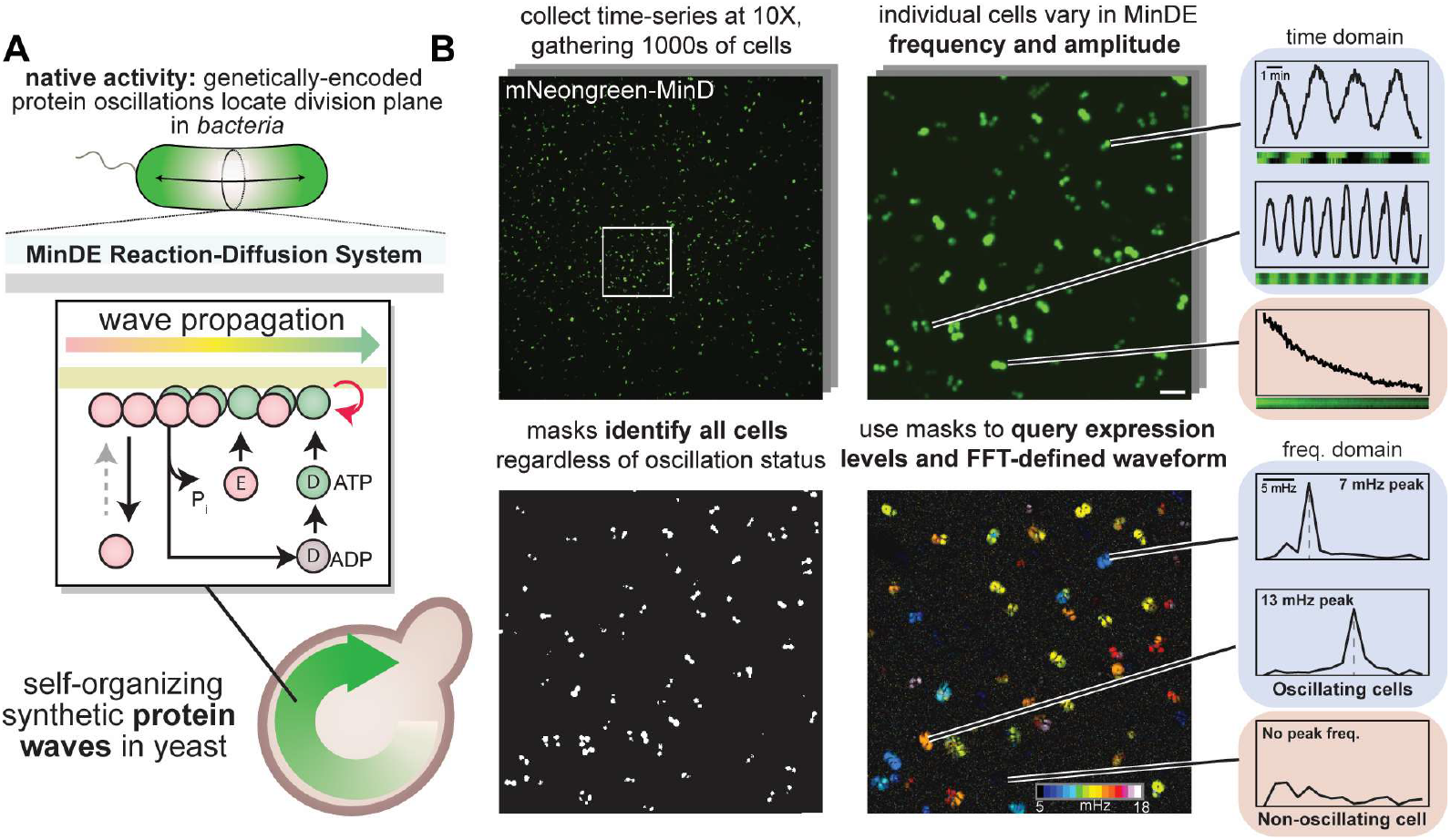
MinDE-family genetically encoded protein oscillations (GEOs) in budding yeast. **A**. MinDE-family ATPase/activator pairs drive protein oscillations based on reaction-diffusion dynamics. MinD-family ATPases dimerize and binds the membrane in their ATP-bound state. MinE-family Activators catalyze the hydrolysis of ATP to drive components off the membrane. When the *E. coli* MinDE genes are expressed in yeast, circular waves form. **B**. Each pixel of a cell has a temporal oscillation, so cells can be imaged at low magnification and high sample size. Cells are masked by fluorescence, and the time trace of each cell’s pixels is then processed with FFT to decode the underlying frequency. Non-oscillating cells have no discernable peak and are not visible on the frequency-colored image. Scalebar = 20 microns.

We first expressed fluorescently tagged MinD ATPase and MinE Activator *E. coli* GEO components on separate Cen/ars plasmids. This expression strategy generates a wide range of ATPase and Activator expression levels that are uncoupled from one another on the population level, providing access to many different configurations of the GEO circuit. Imaging this population by timelapse fluorescence microscopy revealed a large proportion of oscillating cells, in which the GEO waves manifest as a “washing machine” phenomena circling the cell interior and developing bud (Video S1). Within this Cen/ars population, we observed considerable diversity in the waveforms across different expression levels: some cells oscillated slowly (7 mHz), some oscillated faster (13 mHz), and some cells did not oscillate at all (Fig. 1B). Thus, GEOs can realize a wide spectrum of waveforms in budding yeast.

To define the relationship between GEO expression levels and oscillation waveform, we took advantage of the fact that single cell oscillations could be robustly detected at the pixel level even at low (10x) magnification using Fast Fourier Transform (FFT).^31^ This allows the oscillation status (yes/no), waveform (frequency and amplitude) and associated ATPase and Activator expression levels (using fluorescence as a proxy) to be quantitatively analyzed for thousands of individual cells at once from a single field of view. To increase our coverage of the theoretically accessible expression space, we aggregated data from many fields of view for all pairwise combinations of high (2micron) and low (Cen/ars) copy expression plasmids, producing a comprehensive dataset consisting of more than 3500 cells (Fig. S3A). All cell expression data was background-corrected by image.

From these single-cell data, we plotted phase portraits that map ATPase and Activator expression levels to the oscillation state (Fig. 2A), frequency (Fig. 2B) and amplitude (Fig. 2C). The structure of these portraits reveals the key programming logic controlling GEO waveform. Oscillations can only occur within a specific range of Activator:ATPase ratios. To quantify the boundaries between the functional and non-functional space, we developed and applied a machine-learning classifier based on two logistic regression models^34^ to recover the minimum and maximum Activator:ATPase ratios that can support oscillations (Fig. S1). These parameters establish the GEO’s functional frequency bandwidth and the upper and lower limits of its dynamics (Fig. 2A). The model was highly accurate, precise, and sensitive in classifying cells as oscillating (accuracy: 0.938, precision: 0.944, sensitivity: 0.988; detailed in supplement).

**Figure 2:**
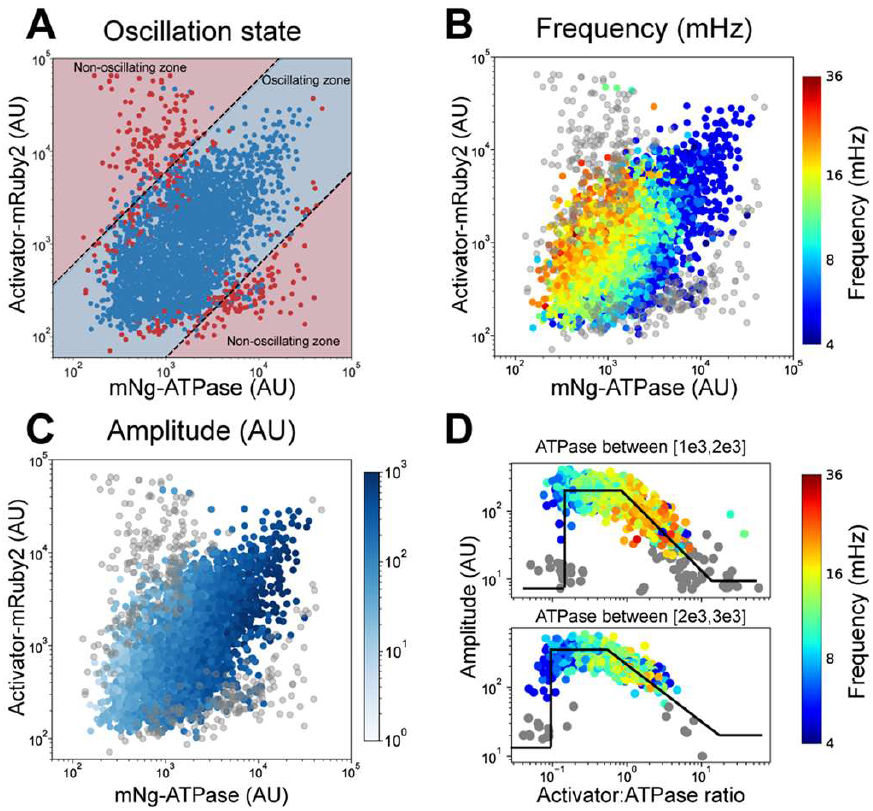
Phase portrait analysis defines how ATPase and Activator expression controls GEO oscillation status, frequency, and amplitude. Budding yeast single-cell GEO behavior, waveform metrics and component expression levels were aggregated from uncoupled plasmid expression data for visualization. **A**. Oscillating cells fell between two Activator:ATPase ratios, which were classified with logistic regressions. The point color shows true status, while the background shading denotes ML-model predicted status. Blue= oscillating, red = non-oscillating. **B**. GEO frequency within the phase portrait. Frequency largely scales with Activator:ATPase ratio. Grey cells are non-oscillating. **C**. GEO amplitude within the phase portrait. Amplitude largely scales with ATPase abundance. **D**. Within vertical slices of the phase portrait, amplitude exhibits three distinct states: non-oscillating, stationary, and decay as expression approaches the upper boundary of oscillation. Frequency scales with Activator:ATPase ratio. N = 3643 cells.

Within the functional oscillation space, GEO frequency increases with increasing Activator:ATPase ratio and GEO amplitude scales with ATPase expression but drops off as the Activator levels approached the boundaries of the functional space (Fig. 2D). For a given slice of ATPase expression, increasing Activator toggles amplitude from background levels to an “on state”, which is maintained until a decay back to background.

Our phase portrait characterization agrees with previous observations of GEO behavior in mammalian cells, *in vitro* systems and mathematical models but provides new experimental clarity over the boundary and transition to non-oscillating states. Taken together, the GEO phase portrait provides a comprehensive description of oscillation behavior and its underlying expression constraints: where oscillations can arise, and how frequency and amplitude change as a function of ATPase and Activator concentration.

### Independent effects of intrinsic and extrinsic expression noise on GEO frequency and amplitude

The GEO phase portraits we constructed in Fig. 2 provide the ATPase and Activator expression levels required to realize a specific oscillation waveform. However, genetically identical cells will vary in component levels owing to expression noise, making it unclear how precisely a specific waveform can be genetically encoded in practice. This noise arises from two separable components: extrinsic noise, which reflects variation between cells in a population; and intrinsic noise, which reflects variation between equivalent promoters within the same cell.^19^

To experimentally determine the effects of intrinsic and extrinsic expression noise on GEO waveform, we generated a yeast strain harboring genomically integrated constructs that separately express each of the *E. coli* GEO ATPase and Activator components from a strong Tef1 promoter. We imaged the dynamics of this genomically-locked strain (henceforth: int-GEO1) using timelapse fluorescence microscopy and applied our analysis pipeline to quantify the resulting expression levels and waveforms present in the population. Compared to Cen/ars, the observed GEO frequency within the int-GEO1 population was remarkably consistent (Fig. 3A, Video S2), with the inner 80% of the population (10^th^-90^th^ percentiles) oscillating between 13 and 17 mHz, as compared to an inner 80% range of 8 to 20 mHz for the uncoupled Cen/ars population. Importantly, the stability of oscillations in the int-GEO1 strain persisted even as cells grew and expanded out over multiple generations (Fig. 3B).

**Figure 3:**
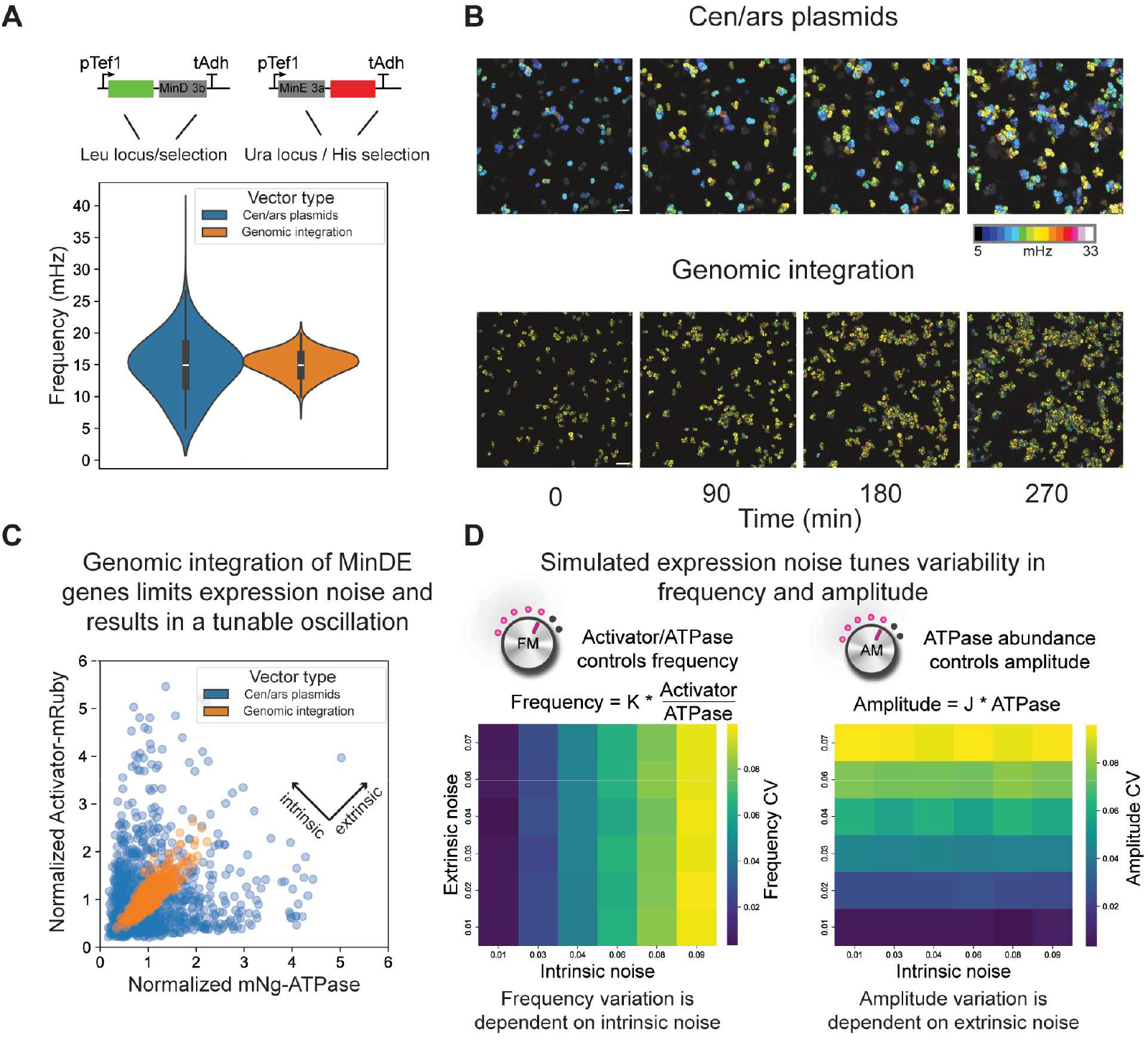
Genomic integration reveals separable effects of intrinsic and extrinsic expression noise on GEO frequency and amplitude. **A**. Integrating the GEO circuit reduced frequency bandwidth compared to the Cen/ars paradigm under the same promoters (N > 1200 for each population). **B**. Integrated GEOs remain robust and predictable during culture expansion, while Cen/ars populations shuffle due to random plasmid copy number inheritance. Scalebar = 20 microns. **C**. Expression data from panel **A** plotted normalized by each population’s means along the given axes. Genomic integration has lower extrinsic and intrinsic noise compared to the Cen/ars population. **D**. Simulated population structures with varying noise show that frequency noise is exclusively dependent on intrinsic noise, while amplitude noise is exclusively dependent on extrinsic noise. CV = Coefficient of Variation.

To clarify the origin of this robustness, we projected the genomically-locked single-cell int-GEO1 data onto the full GEO Cen/ars phase portrait, normalizing by separate population means along both axes (Fig. 3C). This revealed that expression levels of the ATPase and Activator were highly correlated, falling in a narrow band along a fixed Activator:ATPase ratio and consequently a specific GEO frequency. Thus, although extrinsic noise can introduce large variation in component expression levels, these differences are preserved across similar genes and promoters, resulting in low Activator:ATPase ratio variation.

To formalize our understanding of these experimental observations and gain intuition, we simulated waveforms for clonal populations facing different levels of intrinsic and extrinsic expression noise. Briefly, two random normal distributions were used to create expression profiles: the first for ATPase abundance, and the second for a set of “offsets” from the ATPase abundance to generate the Activator expression (Fig. S2A). The result was a set of expression profiles with varying intrinsic and extrinsic noise (Fig. S2B). Tuning the standard deviation of the ATPase distribution affected extrinsic noise, while the standard deviation of the offsets tuned intrinsic noise (Fig. S2C). Guided by the structure of the GEO phase portrait (Fig. 2), we modeled frequency as a linear scaling of Activator:ATPase ratio, and amplitude as a linear scaling of ATPase abundance. We then sampled ATPase and Activator expression levels from the simulated distributions and quantified the resulting frequency and amplitude variation within the resulting population. These simulations confirmed that a consequence of the GEO phase portrait is that frequency and amplitude variation are generally independent of one another, with frequency variation influenced by intrinsic noise and amplitude variation influenced by extrinsic noise.

### Exploiting noise insights to engineer GEO waveform structure within a population

Our analysis has so far revealed that 1) GEOs can generate a diversity of waveforms across different expression levels programmed by an underlying phase portrait; and 2) variation in GEO frequency and amplitude are independently controlled by intrinsic and extrinsic expression noise respectively. As a result, we hypothesized that different GEO circuit expression paradigms that introduce or reduce different sources of noise would enable construction of different clonal populations that robustly oscillate at unique frequencies, or waveform libraries that independently vary frequency or amplitude for barcoding or dynamic perturbation applications.

To develop different noise-guided GEO circuit expression strategies, we domesticated the *E. coli* GEO ATPase and Activator genes as parts for use in the MoClo modular golden gate cloning system.^33^ This allowed GEO components to be easily connected to different promoters to build diverse transcriptional units (Fig. 4A). These transcriptional units can be further combined in different ways to introduce copy-number variation or drive coupled or uncoupled component expression within a population, thus tuning the effective intrinsic or extrinsic expression noise.

**Figure 4:**
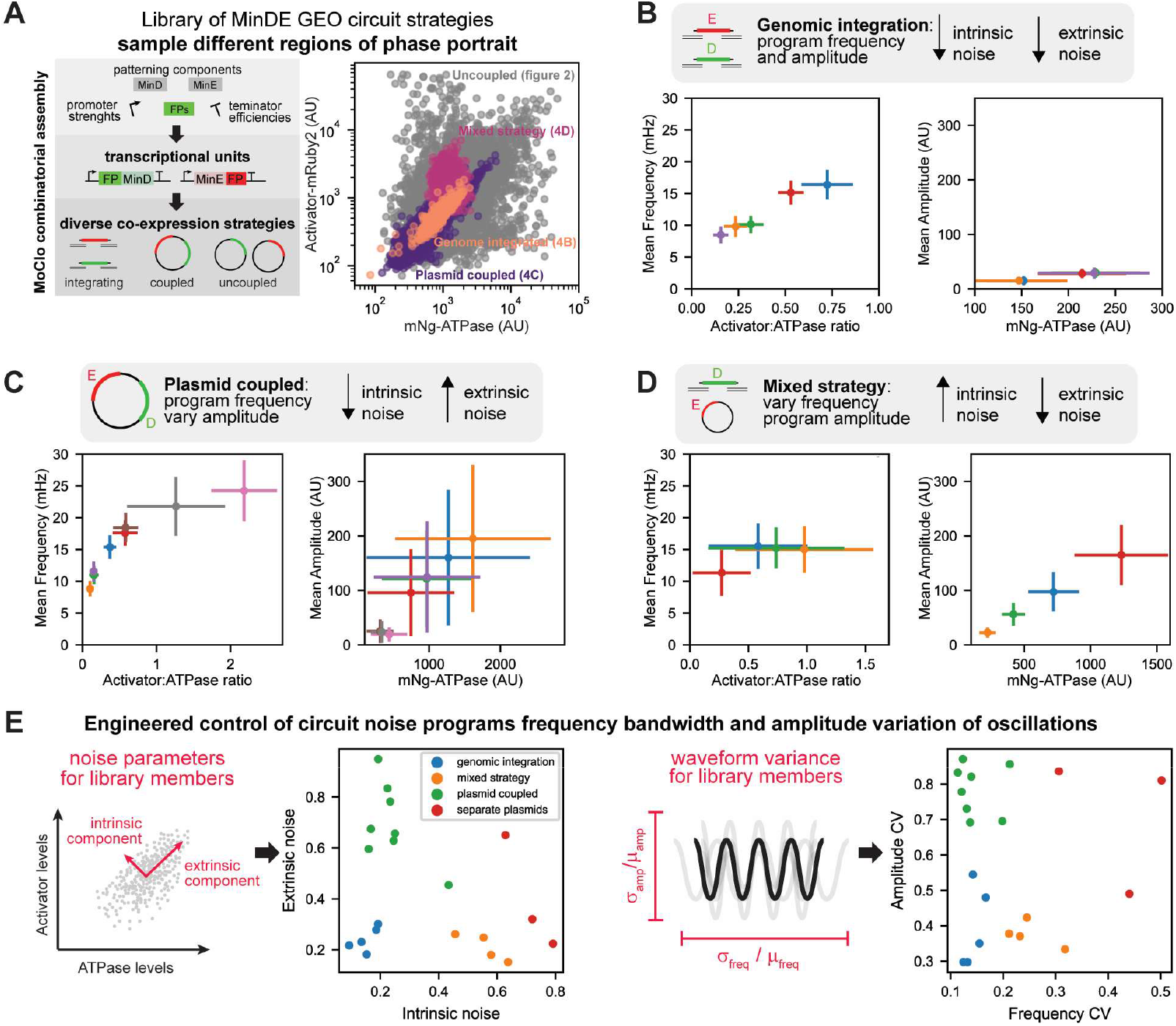
Noise-guided expression strategies for tuning GEO waveform. **A**. GEO ATPase and activator components were domesticated for MoClo toolkit logic, allowing for multipart assembly of transcriptional units (TU) and different co-expression strategies. Scatterplot shows example population distributions for specified expression strategies. **B**. Genomic integration results in frequency and amplitude tunable populations. The means and standard deviations are shown. Point color corresponds to promotor combinations for the ATPase and Activator components. **C**. Expressing GEO activator and ATPase on one plasmid results in populations with low frequency noise, but high amplitude variation. **D**. A mixed expression strategy with integrated ATPase and plasmid Activator results in frequency-varied, amplitude-locked populations. For specific promoter combinations disambiguation for each panel, see Fig. S3. **E**. Population-level noise depends on expression methods, with distinct clusters forming along the intrinsic and extrinsic axes depending on the expression strategy. Expression noise translates directly to GEO oscillation noise in amplitude and frequency. CV = Coefficient of Variation.

Because the int-GEO1 strain produced a tight, near uniform frequency at the population level, we reasoned that genomically integrated populations with different Activator:ATPase ratios would access different frequencies. To this end, we generated ATPase and Activator transcriptional units driven by a range of different promoter strengths and used single-copy integration to deliver these into the same two genomic locations for comparison (Fig.4B, Fig. S3B). In this context, different promoter combinations reliably established different Activator:ATPase ratios within the population, resulting in specific frequencies locked with population standard deviations of ∼2 mHz (Fig. 4B). Amplitude variation was also low, due to the lack of DNA-level variation in ATPase gene copy number.

Seeing that different GEO frequencies can effectively be locked by different genomically integrated promoter combinations, we next sought to create population structures that maintained a consistent frequency but introduced large variation in wave amplitude. To do this, we expressed both the ATPase and Activator transcriptional units with specific promoter combinations on single Cen/ars plasmids. Like their genomically integrated counterparts, these single-plasmid expression strategies showed tight Activator:ATPase ratios (low intrinsic noise) within the population and thus a narrow band of GEO frequencies (Fig. 4C). However, because the copy number of these plasmids varied among individuals within the clonal population, there was extensive variation in total expression and amplitude due to high extrinsic noise (Fig. 4C, Fig. S3C). Thus, coupled expression strategies – whether realized through variable-copy plasmids or genomic integration – serve as effective frequency-locking strategies by maintaining consistent expression ratios even in the presence of extrinsic noise. Because of this immunity to extrinsic noise, as a signal, the frequency varies less on a population level than the fluorescence of the ATPase channel, regardless of expression strategy (Fig. S3F).

Finally, we aimed to create population structures that introduced variation in GEO frequency while maintaining a consistent GEO amplitude. We accomplished this using a mixed strategy design in which ATPase expression was locked by genomic integration, but Activator was expressed from a Cen/ars plasmid. This expression approach aimed to increase population variation in the Activator:ATPase ratio, scrambling GEO frequency, while maintaining a relatively fixed ATPase level, locking in GEO amplitude. Using this strategy with different promoter combinations and number of ATPase genome integrations, we could effectively establish different populations with similar, broad GEO frequency bandwidths but whose oscillations were centered about different amplitudes with low variation (Fig. 4D, Fig. S3D).

We can directly compare these different GEO circuit expression strategies in the context of noise by decomposing their expression variation into intrinsic and extrinsic components. Genomically integrated GEOs had the lowest intrinsic and extrinsic noise, while expressing components from separate plasmids produced the highest intrinsic and extrinsic noise (Fig. 4E). As intended, our plasmid-coupled expression designs had low intrinsic noise but high extrinsic noise; and our mixed expression strategies had low extrinsic noise but high intrinsic noise. Critically, these differences in how expression noise was partitioned directly corresponded with the degree of amplitude or frequency variation observed within a population (Fig. 4E). Taken together, we were able to exploit noise effects to design different GEO expression strategies that establish a spectrum of waveforms with independent tunability over frequency and amplitude variation on the population level.

### Evolutionarily diverse ATPase and Activator modules provide a range of phase portraits for noise-guided encoding of GEO oscillations

The noise-guided waveform encoding strategies we demonstrated for the *E. coli* GEO system in budding yeast operate at the level of gene expression, acting like filters on the GEO phase portrait that isolate specific subsets of expression space. As a result, the waveforms accessible and programmable by these approaches are still constrained by the biochemistry of the GEO ATPase and Activator components that underlie that phase portrait. An expanded set of alternative GEO ATPase/Activator pairs would provide additional phase portraits as starting points for application of our GEO expression strategies in waveform design.

To this end, we isolated and cloned 13 evolutionarily diverse MinDE-family ATPase and Activator modules^32^ into separate Cen/ars plasmids and used a yeast mating strategy to generate all 169 pairwise GEO combinations (Fig. 5A). These constructs expressed MinD ATPase variants N-terminally tagged with mCherry, and MinE Activator variants C-terminally tagged with EGFP. We imaged each system and defined its associated phase portrait using our analysis pipeline. Of the 169 combinations, 66 had >5% oscillating cells, which we defined as the threshold for considering a GEO as functional. From this library we identified new and useful oscillation properties for many of these GEO pairs, where identical expression levels program different frequencies (Fig. 5B, Video S3). To compactly visualize the diversity of GEO behaviors available, we plotted the mean frequency and amplitude for all functional members (Fig. 5C). This allows ready identification of GEO systems that oscillate significantly faster (*P. damselae* MinDE, 44 mHz), or show significantly greater amplitude (*P. poae* MinD, *Dickeya dadantii* MinE, 47 AU) than the original *E. coli* GEO system (mean frequency 12 mHz; mean amplitude 18 AU). Hybrid pairings can also be functional, with the Activator module from fast species boosting oscillations of slower ATPase modules (*E. coli* MinD, *P. damselae* MinE, mean frequency 30 mHz).

**Figure 5:**
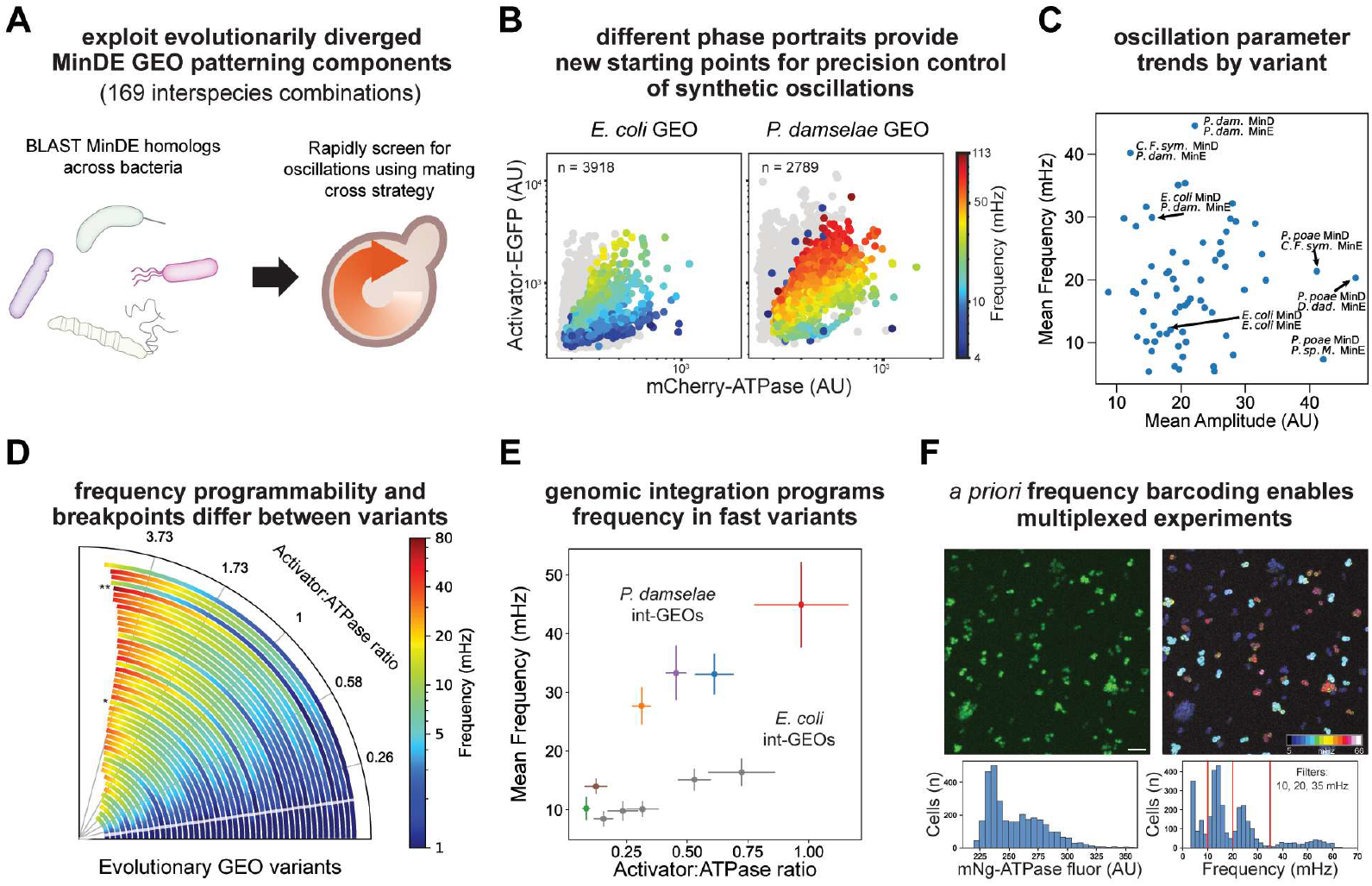
Evolutionary diversity of GEO components uncovers new phase portraits for noise-guided waveform design and application. **A**. 13 MinD-family ATPases and MinE-family Activators from across bacteria were cloned for uncoupled plasmid expression and tested using a mating cross to generate 169 GEO combinations. **B**. Different GEOs had vastly different phase portraits, with the same expression levels resulting in markedly different frequencies and amplitudes. Gray cells are non-oscillating. **C**. GEO phase portraits had diverse mean frequencies and amplitudes, with the *P. damselae* native pair having the highest mean frequency, and the *P. poae* MinD + *D. dadantii* MinE having the highest mean amplitude. **D**. Radial “speedometer” plot of modeled fit parameters, visualizing the range of Activator:ATPase ratios that support oscillation colored by frequency for all well-fit GEO library members side-by-side. The white line denotes the existence of a minimum Activator:ATPase ratio. * = *E. coli* GEO; ** = *P. damselae* GEO. **E**. Genomic integration of the fast *P. damselae* MinDE proteins creates fast and tunable clonal populations. The same promoter combinations for *E. coli* GEO are shown in grey (see figure 4B). Error bars denote standard deviation. **F**. Iso-frequency populations enable single-fluorophore barcoding of clonal populations. These cells exist in similar expression regimes, but distinct frequencies, permitting multiplexed experiments on pooled populations barcoded by frequency. Here, four populations are decoded with frequency filters at 10, 20, and 35 mHz. Histograms represent distribution of fluorescence and frequency in entire field of view. Scalebar = 20 microns.

To further analyze the phase portrait structure of each GEO library member, we used our ML-classifier to define the upper limit on Activator:ATPase ratio for each GEO’s functional oscillation space. These models returned good classifications (see supplement) and upper boundary slopes ranging from 1.25 to 10.67 depending on GEO. We then modeled how frequency is programmed within the functional space by fitting frequency to a linear regression of the Activator:ATPase ratio (Fig. S4A). The metrics for all functional and well-fit GEO pairs can be visualized side-by-side using a radial “speedometer plot”, which shows the range of ratios that support oscillation for each GEO pair along the radial axis color-coded by the frequency they generate at that ratio (Fig. 5D). Here, the same Activator: ATPase fluorescence ratio resulted in markedly different results depending on the GEO components, both in oscillation state and frequency. This toolbox of phase portraits suggests new waveform programming strategies that take advantage of differences in the binary state of oscillation and frequency bandwidth of different GEO systems. Some GEOs have a wide operable expression range, offering many subsets of expression for frequency locking. Others, like the *P. damselae* ATPase + *P. sp. Mhis* Activator, scale frequency rapidly (Fig. S4A) but have a low upper bound, resulting in a sensitive but narrow operating range.

To demonstrate how this expanded collection of GEO phase portraits can be combined with the precision genetic control strategies we developed earlier, we analyzed the oscillation behavior of the fast and robust *P. damselae* GEO (mean 44 mHz) using the same promoter combinations we previously analyzed for the *E. coli* GEO. Since we domesticated these parts in our toolkit, we returned to the original FP designation of mNeongreen-MinD and MinE-mRuby2. While the expression levels of the GEO components are similar between matched promoter combinations (Fig. S3B,E), the different GEOs give rise to clonal populations that oscillate at unique, well-resolved frequencies (Fig. 5E). We used *E. coli* and *P. damselae* int-GEOs to spectrally barcode four separate clonal populations and deconvolute clone identity in mixtures based solely on single-cell frequency in a single fluorescent channel (Fig. 5F).

The diverse portfolio of GEO systems we characterized provide a rich assortment of phase portraits for noise-informed waveform programming. By coordinating differences in component biochemistry with the different GEO circuit expression strategies we developed earlier, we expand the space of oscillations that can be precisely encoded within a population.

## Discussion

In this work, we demonstrate noise-guided control strategies for genetically encodable oscillators (GEOs) that leverage evolutionary diversity of GEO components and different expression methods within the budding yeast model organism. Key engineering insights came from the construction and quantitative analysis of GEO phase portraits, which reveal the spectrum of waveforms accessible to a given GEO circuit and how ATPase and Activator expression levels, noise, and biochemical features define oscillation status and waveform properties. These phase portraits allowed us to model and experimentally define how different sources of expression noise independently control the variation in a GEO’s frequency and amplitude. Using this knowledge, we developed a suite of GEO expression strategies that act like subtractive filters on the broad “white noise”-like distribution of waveforms within a GEO’s phase portrait.

GEO frequency is ratiometrically encoded in the relative amounts of ATPase and Activator, making it naturally buffered against extrinsic expression noise. Target waveforms could thus be reliably encoded using single copy integration circuits that combine different promoter strengths with GEO components from different species. Clonal populations could be barcoded by frequency within a single fluorescent channel, allowing multiplexed phenotypic analysis or interrogation of the spatial structure of populations of mixed-cell types or lineages. A dedicated biochemical carrier signal can thus be hard-wired in a cell’s genome for AM or FM encoding of dynamic single cell data using more elaborate circuit designs.^23^ Not only does frequency provide an added dimension of data encoding, but the time-resolved unit (mHz) is concrete and transferable compared to the arbitrary units (AU) of fluorescence alone. Constructs that encode specific GEO waveforms could potentially be pooled and DNA-barcoded for use in sequencing-based readouts that relate specific dynamics to phenotypic outcomes.

Alternatively, we could harness the separable effects of intrinsic and extrinsic noise on GEO waveform frequency and amplitude noise. Inherently shuffled amplitude libraries can be generated within specific frequencies (using coupled, single plasmid expression), or diverging frequency libraries can be standardized within specific amplitude ranges (using mixed, uncoupled expression). These expression paradigms could potentially be combined with additional strategies that operate downstream of gene expression to further buffer against noise, such as feedback regulation of transcription,^35^ protein release from condensates,^36,37^ or cell cycle synchronization.^38^ Conversely, a genome-integrated GEO could provide a dynamic reporter of expression noise, using the frequency and amplitude variation as a single-fluorophore population reporter of noise. By using different methods of expressing GEO components, waveforms can be generated and applied to a diverse range of experimental needs, not only in budding yeast but also other eukaryotic systems.

These engineering strategies emulate how natural systems exploit noise as a tunable parameter – quenching it when precision is needed or exploiting it to introduce population-level variation during stress or division of labor.^39–41^ While our study focused on oscillatory dynamics, we anticipate that noise-guided circuit design principles will be valuable for understanding and engineering dynamic protein circuits more generally. For instance, synthetic networks of interacting kinase/phosphatases,^42^ GEF/GAPs,^43,44^ receptors/ligands,^15,45,46^ or protease/substrates^14,47,48^ have recently been established as tools for dynamic protein circuit design. Because expression level effects and component ratios will impact the responsiveness and timescales of these circuits, aligning gene expression and noise effects with dynamic outputs would improve robustness and performance. Our work highlights the value of considering the impact and effects of noise in protein circuit dynamics, not as something detrimental to their performance but as a powerful tool for distributing different dynamics within a population.

**Acknowledgements**.

We thank members of the Coyle Lab, A. Weeks, M. McClean and W. Bement for advice, helpful discussions, and critical reading of the manuscript. We thank M. McClean, A. Gasch, and E. Craig for assistance and sharing of yeast strains, vectors, and application of the MoClo kit.

## Author contributions

DB and SMC conceived the overall project. The experimental plan was implemented by DB, EWZW, and TG. DB collected all experimental data for the paper with assistance from EWZW and TG. EWZW collected most of the data in the GEO variant screen (Fig. 5A-D). DB developed all analysis pipelines apart from the logistic regression models for oscillation state classification, which were developed by EWZW. RR and TG contributed to the selection of ATPase and Activator GEO variants tested in Fig. 5 and provided helpful feedback around frequency linear modeling. DB and SMC prepared the figures and wrote the manuscript with suggestions from all authors. SMC supervised all aspects of the work.

