## Supplemental Methods and Materials for "Noise-guided tuning of synthetic protein waves in living cells"

**Affiliations:**
[1]Department of Biochemistry
[2]Cellular and Molecular Biology Graduate Program
[3]Integrated Program in Biochemistry Graduate Program

University of Wisconsin-Madison, Madison, Wisconsin 53706, USA.

### Methods

#### Materials.

Table S1: Materials used in this study

Table S2: Yeast strains generated in this study

Table S3: GEO species MinDE sequences and abbreviation disambiguation

*S. cerevisiae* strains BY4741 (*MAT***a** *his3*Δ1 *leu2*Δ0 *met15*Δ0 *ura3*Δ0) and BY4742 (*MAT***α** his3Δ1 *leu2*Δ0 *lys2*Δ0 *ura3*Δ0) were obtained from the Craig and Weeks lab at UW-Madison.

#### Gene synthesis and cloning.

The modular yeast toolkit[1] was obtained from the McClean lab at UW-Madison. Further part plasmids were generated using the golden gate procedures described by Lee et. al. (2015) and NEB. The ymNeongreen fluorescent marker was cloned from addgene #125704 from Botman et. al (2019)[2] using a PCR product with forward primer 5' - GCATCGTCTCATCGGTCTCATATGGTATCTAAGGGTGAAGAA -3' and reverse primer 5' – ATGCCGTCTCAGGTCTCAAGAACCCTTGTACAATTCGTCCATACCC – 3' which introduces a point silent mutation at the start of the protein to allow compatibility with the Moclo workflow. All remaining MinD, MinE proteins were cloned or ordered from Twist or IDT following the same cloning logic, with flexible linkers at the N or C termini, respectively. MinD genes were cloned as parts 3b and MinE as parts 3a. 3b MinD parts contained a stop codon to keep the C-terminus membrane-targeting sequence unobstructed by a GS linker. DNA was prepared from KCM transformation into DH5alpha or XL10-Gold cells following overnight growth and miniprep with Qiagen kits.

#### Yeast culturing, transformation, and imaging methods.

Budding yeast were transformed with plasmids and integration vectors following a standard LiOAc + ssDNA method[3]. YPD was used for preparing cells for transformation, which contains 1% (w/v) Bacto Yeast Extract, 2% (w/v) Bacto Peptone, 2% (w/v) Dextrose. Transformation products were plated on auxotrophic selection agar plates containing 0.67% (w/v) Yeast Nitrogen Base Without Amino Acids (Sigma #Y0626), 2% (w/v) Bacto Agar, 2% (w/v) Dextrose, and 0.2% appropriate amino acid supplement (with specific auxotrophic selection). The individual amino acids concentrations are defined in Yeast Synthetic Drop-out Media Supplements (Sigma-Aldrich, Saint Louis, Missouri).

**Yeast GEO variant screen.** We constructed Cen/ars plasmids expressing MinD and MinE variants from the mammalian vectors in Rajasekaran et al. (2025).[4] Briefly, the mCherry-MinD and MinE-EGFP genes were isolated with restriction digestion (BamHI and NotI), then ligation cloned into custom Cen/ars vectors with the pTef1 promoter and Adh1 terminator flanked by BsaI cut sites compatible with the overhangs of the genes. The MinD plasmids had the leucine selection marker, and the MinE plasmids had the histidine selection marker. We transformed the MinD and MinE plasmids into *S. cerevisiae* strains BY4741 (*MAT***a** *his3*Δ1 *leu2*Δ0 *met15*Δ0 *ura3*Δ0) and BY4742 (*MAT*α his3Δ1 *leu2*Δ0 *lys2*Δ0 *ura3*Δ0)[5] respectively using the usual LiOAc/ssDNA transformation protocol[3]. Single colonies were streaked in stripes on the appropriate selection media plates, then a mating cross was performed by replica-plating these stripes first onto YPD, growing them overnight, then onto double-selection plates (-his -leu) to select for diploid cells carrying both plasmids.

**Imaging preparation.** Low fluorescence media (LFM) was made consisting of 0.17% Yeast Nitrogen Base without Ammonium Sulfate, Folic Acid, or Riboflavin (#MP114030512, Thermo Fisher Scientific, Waltham, Massachusetts), 2% Dextrose, 0.5% Ammonium Sulfate, 0.2% appropriate amino acid supplement (with specific auxotrophic selection). The individual amino acids concentrations are defined in Yeast Synthetic Drop-out Media Supplements (Sigma-Aldrich, Saint Louis, Missouri).

Single colonies of yeast were inoculated in 1mL appropriate auxotrophic LFM in Nunc™ 96-Well Polypropylene DeepWell™ plates (Thermofisher #95040462) and grown overnight in a shaking incubator (180RPM) at 30 ºC. The following morning OD600 values were measured, and cells were diluted to 0.01 OD, then allowed to grow for 4-6 hours. A 96-well glass plate was prepared by pipetting 100 µL ConA (2mg/mL) into each well and was incubated for 1 minute, then removed. Yeast cultures were transferred to 1.5 mL Eppendorf tubes and vortexed vigorously. 200 µL of yeast culture was then added to the well, then removed after 10-15 minutes of settling. 100 µL of fresh LFM media was carefully added on top of the cells.

Cells were imaged with a Tokai stage top incubator. All temperatures were set to 30 ºC except for the top heater, which was set to 37 ºC to prevent condensation.

**Fluorescence microscopy.** Cells were imaged using on a Nikon Ti2-Eclipse. Most data were imaged using the Photometrics 95B 25mm camera at 10X magnification with the Optivar. Data in figure 3C and 5F,G were imaged using the Hamamatsu Orca-fire camera at 10X magnification without the Optivar. Time-series were collected in the green and red channels for 10 minutes with 3 second intervals (2 second interval in the MinDE variants experiments (figure 5)).

**Data analysis and modeling**

**Image processing.** timelapse fluorescence images were processed with custom-written Python scripts. For cell segmentation, the red and green channels were time-averaged and blurred with a gaussian blur. An adaptive threshold was applied to identify fluorescent pixels and create a mask (openCV package).[6] Cells were identified in the mask using scipy ndimage functions.[7] Fluorescence, size, and location data were extracted using numpy,[8] and frequency information was extracted as previously described.[9] Cells were filtered from noise using size filters. Binary oscillation state was determined using filters on pixel-level frequency variance and corrected amplitude (amplitude / mean MinD fluorescence) filters (supplement). Frequency colored images were made using the real component of the FFT image stack using the temporal color code function in Fiji.[10]

**Logistic regression models.** Analysis was performed using custom-written Python scripts. A median ratio of MinE to MinD was calculated per hybrid pair using green and red fluorescence within the oscillating class. Cells were labeled as above or below the median and as oscillating or not. The data was then split and stratified by these two classes into a training set, reserving one third of the data for testing. Two logistic regression models were then instantiated using the clogistic Python library.[11] One logistic regression model was trained on cells above the median ED ratio to categorize them as oscillating or non-oscillating. The second logistic regression model was then trained on this filtered set in the same manner. Binary predictions from the two models were combined by elementwise multiplication. These predictions were evaluated on the hold-out test set. The decision boundaries produced by each model were also plotted together.

**GEO variant library functional boundary models.** Models were trained as in the previous case, with the following alterations. Oscillating cells that fell below the median MinE:MinD ratio were excluded from analysis, as there were insufficient non-oscillating data for classifying the bottom boundary. A single model was trained for each oscillating MinDE pair, detecting the upper functional MinDE ratio. With this strategy, we were able to create a well-fitting model for each of the 66 oscillating hybrid pairs.

**GEO variant frequency linear regression.** For each frequency bin within a variant, the mean Activator:ATPase ratio was calculated. A 0-intercept linear regression (sklearn.linear_model.LinearRegression) was fit to these mean values with the Activator:ATPase ratio on the x-axis and Frequency in mHz on the y axis (Fig. S4). Bins with fewer than 10 cells were removed from analysis. The slope of these lines was used for generating Fig 5D, in addition to the upper boundary slope for each GEO variant determined by the logistic regressions described above.

**Noise analysis.** Prior to calculating noise, expression was normalized to a standard range by dividing cell expression by the population mean for that sample along the two axes separately. Populations with different expression paradigms were analyzed to determine the extrinsic and intrinsic noise, using the formulas from Elowitz et al.[12] Only the oscillating cells were considered for direct comparison to GEO signal noise. The noise within the frequency and amplitude parameters were calculated by dividing population standard deviation by the population mean.

**Noise simulation.** A standard normal distribution was used to create populations with defined means and standard deviation in one dimension. Another standard normal dimension was used to create a set of offsets of the first. Noise was calculated over a range of standard deviations in the simulated MinD and simulated MinE distributions, as well as a predicted frequency and amplitude noise parameter. (Fig. S2) These results were then binned by intrinsic and extrinsic noise, and the means per bin of the frequency and amplitude noise were plotted (figure 3D).

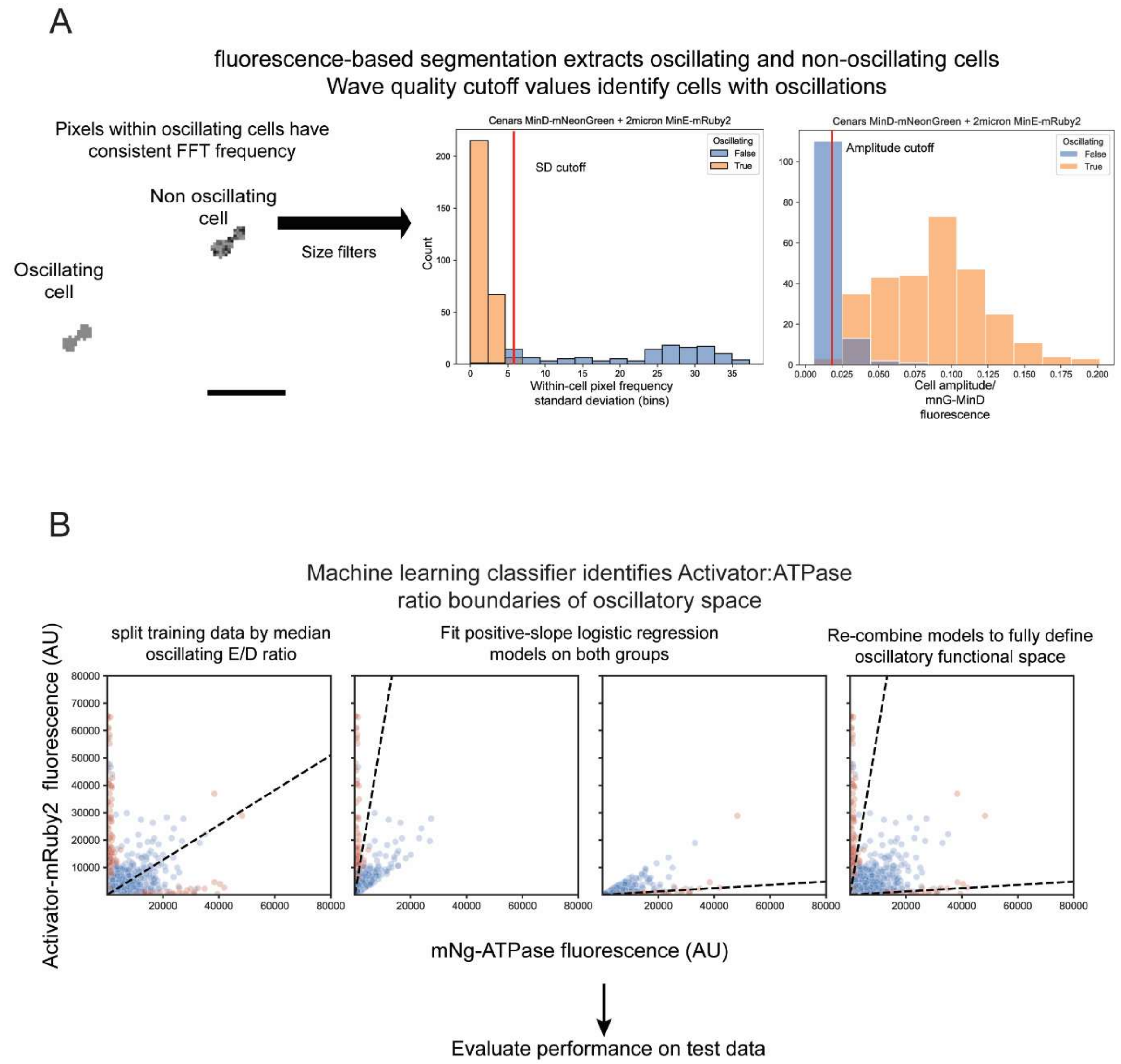


**Figure S1: Imaging and ML approaches to understanding oscillation. A.** We determined cells' oscillation status as a combination of pixel-level frequency consensus within a cell, as well as a minimum amplitude at the peak frequency when normalized against the cell mNeonGreen-MinD fluorescence. Thus, oscillating cells have low variation in frequency between their pixels and high enough amplitude to distinguish a single frequency clearly. **B.** We developed a machine learning classifier to identify the minimum and maximum MinE:MinD ratios that permit oscillation. First, data were split into a training and testing set. Within the training set, two logistic regression models were trained on the upper and lower halves of the data, split by the median Activator:ATPase ratio of the oscillating class. The models were then combined to classify the binary oscillatory state and evaluated on the test set.

For the model shown in figure 2A, the full classification report (on the test set) is as follows:

{'Not Oscillating':

{'precision': 0.8395061728395061, 'recall': 0.5230769230769231, 'f1-score': 0.6445497630331753, 'support': 130.0},

'Oscillating':

{'precision': 0.9447415329768271, 'recall': 0.9878844361602982, 'f1-score': 0.9658314350797267, 'support': 1073.0},

'accuracy': 0.9376558603491272, 'macro avg': {'precision': 0.8921238529081665, 'recall': 0.7554806796186107, 'f1-score': 0.805190599056451, 'support': 1203.0}, 'weighted avg': {'precision': 0.9333694657965678, 'recall': 0.9376558603491272, 'f1-score': 0.9311127174022109, 'support': 1203.0}}

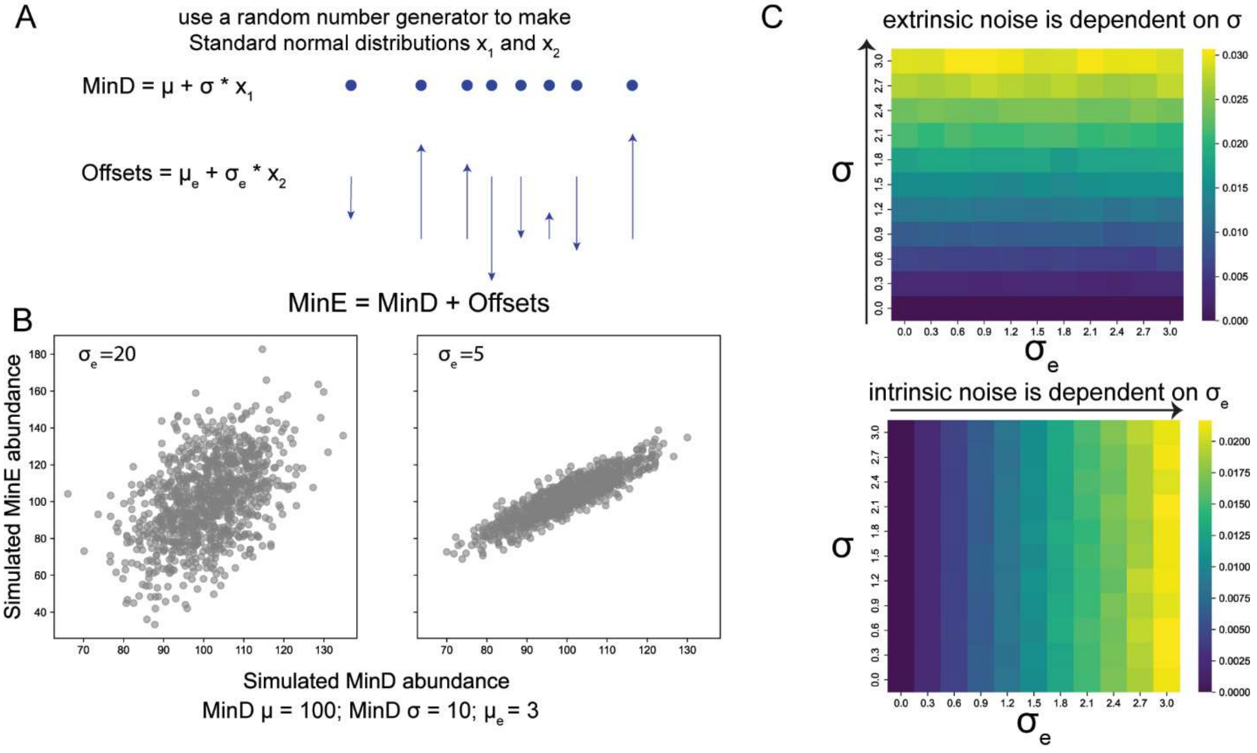


**Figure S2: Simulating expression noise effects on GEO properties. A.** We used two standard normal distributions to simulate MinD abundance and a set of offsets, which when added to the MinD abundance simulates MinE abundance. Thus, we can tune the co-variation of MinE with MinD using the offset set. **B.** Example distributions showing that the intrinsic noise is tunable through the $\sigma_e$ parameter. The means of MinD and MinE are tunable through the $\mu$ and $\mu_e$ parameters, respectively. **C.** A grid simulation iterating over $\sigma$ and $\sigma_e$ values shows that we control the intrinsic and extrinsic noise with these parameters. These simulated populations were then used for the simulated frequency and amplitude distributions in figure 3D.

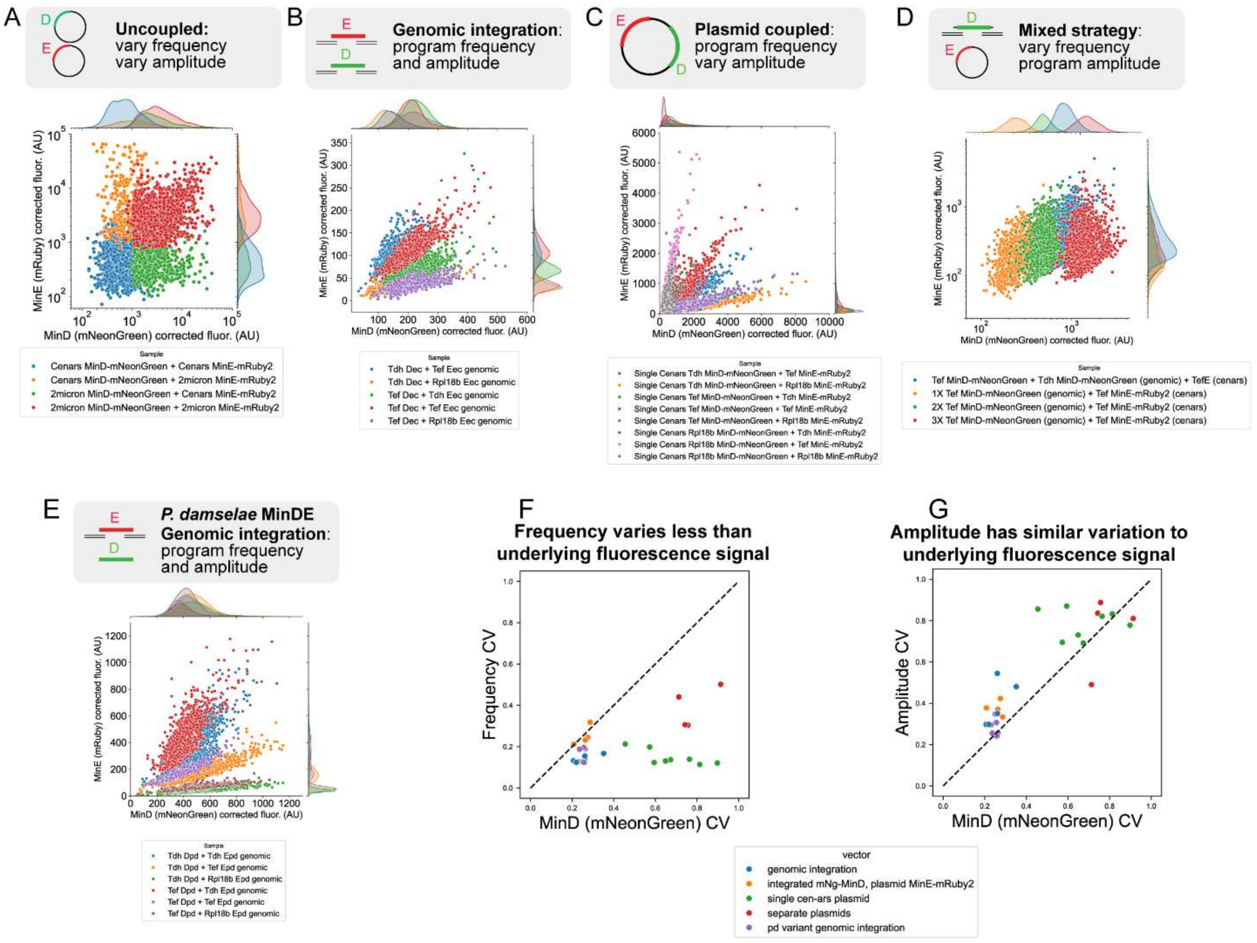


**Figure S3: Raw data showing population structure with different expression strategies. A.** Uncoupled expression with Cen/ars and 2micron plasmids. **B.** Genomic integration of *E. coli* MinDE genes. **C.** Plasmid coupled expression of *E. coli* MinDE genes. **D.** Mixed expression strategy – integrated MinD and Cen/ars expressed MinE from *E. coli*. **E.** Genomic integration of *P. damselae* MinDE genes. **F.** Frequency is a less noisy signal on the population level compared to raw fluorescence. This has to do with the ratiometric programming of this phenomenon. **G.** Amplitude has similar variation to its fluorescence signal, as programmed directly by ATPase abundance.

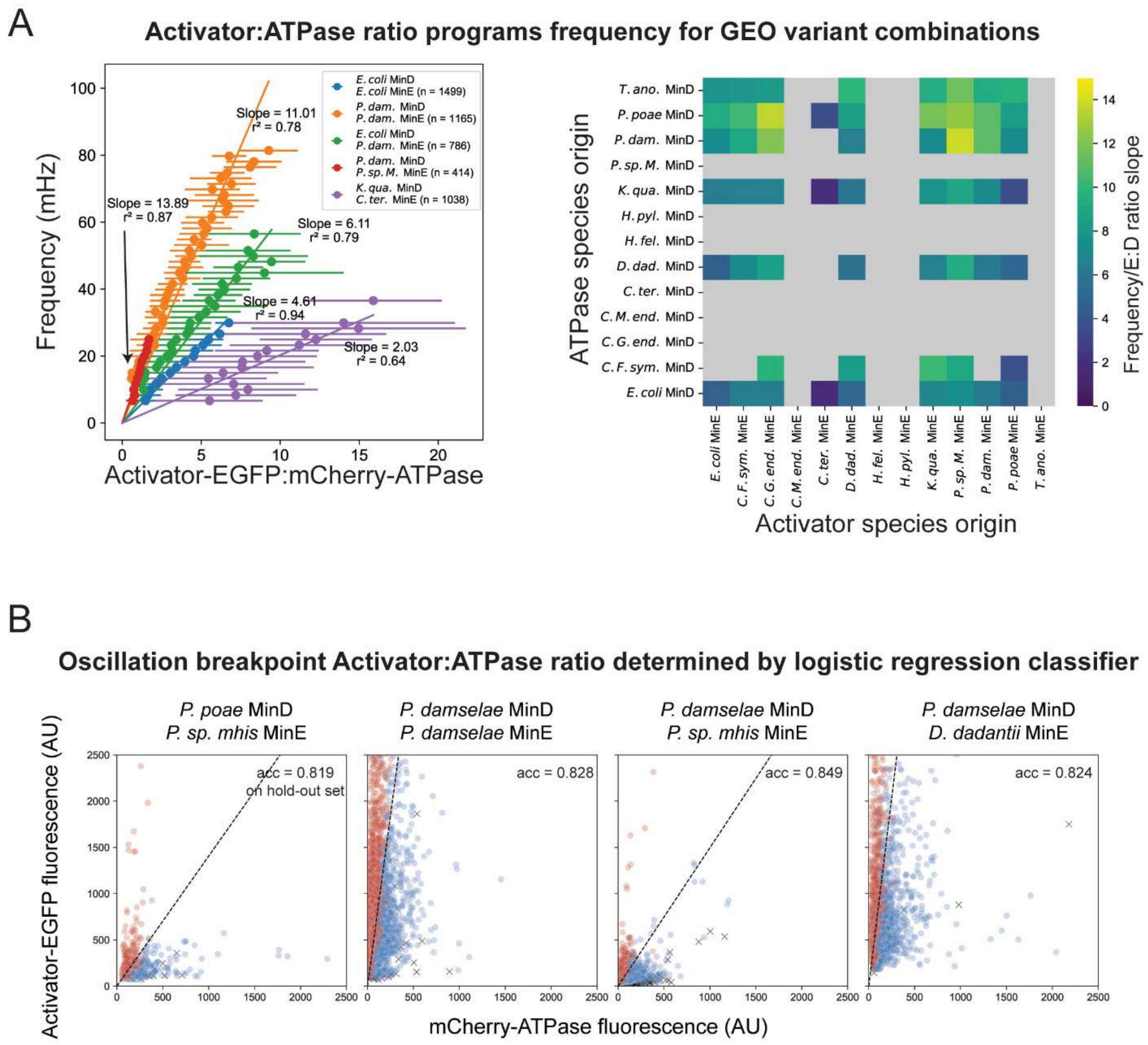


**Figure S4: Quantitative description of GEO variant phase portraits. A.** The Activator:ATPase ratio determined the frequency for each GEO, but the slope of the frequency differed. The highest slope was for *P. dam* MinD + *P. sp. M.* MinE hybrid GEO, although the breakpoint of oscillation was low, meaning that no cells actually reached a high frequency, unlike the *P. dam* MinDE native pair. The lowest frequency slope was observed in the *K. qua* MinD + *C. ter.* MinE hybrid cross. The heatmap shows the frequency to MinE:MinD ratio slopes for 56 of the 66 oscillating GEOs (those with at least 3 summary points to fit a linear regression). Grey squares resemble a cross that did not oscillate in at least 5% of cells and the ten oscillating GEOs not fit. **B.** We applied the logistic regression model described in figure S1 to classify the upper Activator:ATPase boundary of the oscillatory space. This boundary varied greatly between variants. Blue dots are oscillating cells, while red denotes non-oscillating cells. Black crosses represent non-oscillating cells below the median Activator:ATPase ratio, which were omitted for this analysis, as there were insufficient points to classify the lower boundary. The accuracy of classifying cells above the median ratio on the hold-out set is shown for each model. Future work could focus on determining the lower boundary by imaging cells in the region of expression below the median line.

**Data summary tables.**

Table S4: Expression methods summary statistics, noise, etc.

Table S5: GEO variant summary statistics and fit parameters.

**Supplemental videos**

Video S1. Example data from cells with uncoupled Cen/ars MinD and MinE (mNeongreen-MinD shown), showing a zoom from the entire field of view to a small subset of cells. Sped up 150x. The scalebar is 20 microns.

Video S2. Example data from cells with coupled genomic MinD and MinE (mNeongreen-MinD shown), showing a zoom from the entire field of view to a small subset of cells. Sped up 150x.

Video S3. Examples of different ATPase-Activator pairs from the GEO variant screen. mCherry-MinD shown. Scalebar is 25 microns.
